# Bacterial pore-forming toxin pneumolysin drives pathogenicity through shed toxin-loaded host extracellular vesicles

**DOI:** 10.1101/2023.10.12.561978

**Authors:** Saba Parveen, Chinmayi V Bhat, Shaheena Aziz, J Arya, Asmita Dutta, Somit Dutta, Sautan Show, Kuldeep Sharma, John Bernet Johnson, Upendra Nongthomba, Anirban Banerjee, Karthik Subramanian

## Abstract

*Streptococcus pneumoniae* is a global priority respiratory pathogen that kills over a million people annually and produces the pore-forming cytotoxin, pneumolysin (PLY). Host cells expel membrane assembled toxin by shedding microvesicles, but the composition and pathophysiological sequelae of the toxin-induced host extracellular vesicles (EVs) are unknown. Here, we found that EVs shed from PLY-challenged monocytes (PLY-EVs) harbor membrane-bound toxin that induced cytotoxicity upon fusion with recipient cells. EVs from human monocytes challenged with recombinant PLY as well as PLY-expressing pneumococcal strains, but not the isogenic PLY mutant, primed dendritic cells and evoked higher pro-inflammatory cytokines upon infection. Proteomic analysis revealed that PLY-EVs are enriched for key antimicrobial and inflammatory host proteins such as IFI16, NLRC4, PTX3 and MMP9. *In vivo*, zebrafish and mice administered with PLY-EVs showed mortality, pericardial edema, tissue damage and inflammation. Our findings show that host EVs bearing the cytotoxin PLY constitute a previously unexplored mechanism of pneumococcal pathogenesis.

## Introduction

*Streptococcus pneumoniae* (the pneumococcus) causes human upper respiratory tract infections and kills over a million people worldwide annually (*1*). Globally, *S. pneumoniae* is a major causative agent of mortality due to pneumonia in children below 5 years (*2, 3*) and the elderly above 70 years. In 2017, WHO declared *S. pneumoniae* as one of the top 12 priority pathogens that urgently require new antibiotics. *S. pneumoniae* colonizes the nasopharynx of 27–65% of children and <10% of adults asymptomatically (*4*). However, bacterial invasion into the lower respiratory tract and spread to internal organs leads to life-threatening diseases such as pneumonia, septicemia and meningitis. Antibiotic misuse and horizontal transfer of antibiotic-resistance genes have led to the alarming spread of drug resistance amongst pneumococcal strains (*5*). The existing pneumococcal polysaccharide and conjugate vaccines offer limited capsular serotype-dependent protection and have reduced the overall disease burden (*6*), but the emergence of non-vaccine serotypes is a major problem worldwide (*7*).

The pore-forming cytotoxin, pneumolysin (PLY) is a key virulence factor that plays a major role in colonization, invasion and transmission of the pneumococcus (*8*). The binding of PLY monomers to membrane cholesterol triggers protein oligomerization into a 40 to 50-mer prepore structure, that collapses vertically to form β-barrel pores of 30-50 nm in host cells (*9-11*). Pore-formation induces calcium influx into the cell, leading to membrane instability, osmotic imbalance and eventually cell death. However, at sublytic doses of PLY, calcium-dependent membrane repair processes avert cell lysis by plugging the PLY-induced pores and eliminating the membrane pores as microvesicles (*12, 13*). Both microvesicle shedding and endocytosis contribute to removal of PLY-pores from the damaged cell membrane (*14-16*). Extracellular vesicles (EVs) are classified into three major types namely, exosomes (30–200 nm), microvesicles (100–1000 nm), and apoptotic bodies (>1000 nm) based on the size and biogenesis mechanisms (*17, 18*). Exosomes are endocytic in origin, and are released when the intraluminal vesicles of the endosome fuse with the plasma membrane. This is primarily mediated by the endosomal complex required for transport (ESCRT) proteins and Rab GTPases (*19*). Microvesicles are relatively heterogeneous population of vesicles, that are generated by direct outward budding from the plasma membrane (*20*). Susceptibility to PLY-induced cell death varies across cell types, with myeloid cells such as monocytes showing higher resistance as compared to lymphoid cells (*14*). This is due to the preferential binding of PLY to the T cell membrane and the enhanced ability of monocytes to eliminate membrane pores through microvesicle shedding. However, the pathophysiological sequelae of the shed EVs remain unknown. A recent study showed that PLY-induced microvesicles derived from HEK293 cells caused phenotypic changes in macrophages (*21*), but this study was performed using synthetic PLY-bearing liposomes, which do not represent EVs shed from PLY-challenged immune cells. Since EVs transport cargo between cells (*22*), it is intriguing to study the proteomic cargo of EVs shed from immune cells in response to PLY challenge and their implications on host response to infection. In this study, we investigated the fate and downstream consequences of toxin-loaded EVs shed from challenged host cells.

Here, we found that PLY-induced EVs (PLY-EVs) are enriched in innate immune and inflammatory proteins, which induce human dendritic cell (DC) maturation and alert them to infection. *In vivo*, administration of PLY-EVs induced toxicity and inflammation in zebrafish and mouse models. Our findings suggest that expelling PLY toxin pores through vesicles works as double edged sword. While cells get rid of the membrane assembled toxin pores through vesicle shedding to evade lysis, the released EVs containing PLY and other host inflammatory proteins induce cytotoxicity and inflammation in recipient cells. Thus, extracellular vesicles constitute a novel mechanism of dissemination of bacterial toxin pathogenicity.

## Results

### Pneumolysin enhances shedding of extracellular vesicles from monocytes at sublytic doses

Myeloid cells, especially monocytes show greater resistance to PLY-induced lysis in comparison to lymphoid cells, due to their enhanced vesicle shedding capacity. Hence, we used human THP-1 monocytes as a model to investigate the properties and fate of vesicles shed in response to PLY. The conditioned culture supernatant was subjected to differential centrifugation to remove cell debris, apoptotic bodies and larger microvesicles, followed by ultrafiltration to remove non-EV associated proteins. The EVs were subsequently precipitated using ExoQuick-TC reagent (System Biosciences) (**Fig. S1**). THP-1 cells were stimulated with incremental doses of recombinant PLY (0.1-2 μg/ml) for 24h in culture medium pre-depleted of serum EVs, wherein the viability of the treated cells was found to be ∼80% upon treatment with 0.1 and 0.5 μg/ml of PLY (**Fig. S2A, B**). A lytic dose of PLY (2 μg/ml) was used as positive control. Inhibition of Ca^2+^-dependent membrane repair processes using EDTA, significantly reduced the cell viability by ∼50% even at the usual sublytic PLY doses (**Fig. S2C**). To capture the dynamics of PLY-induced EV shedding in real time, we performed high-speed confocal laser scanning microscopy of live THP-1 monocytes, which were labelled with the lipophilic membrane dye, Nile-red and challenged with 0.5 μg/ml of PLY. By ∼14 minutes post stimulation, we found membrane budding and subsequent release of Nile-red positive vesicles from the plasma membrane (**Fig. 1A**, **Video S1**). In contrast, untreated cells released relatively fewer EVs over the same time period **(Fig. 1A**, **Video S2)**. The morphology of the isolated EVs was visualized by negative contrast transmission electron microscopy which showed that the particles were spherical, membrane bound and heterogeneous in size (**Fig. 1B**). Dynamic light scattering analysis showed that EVs purified from untreated cells (naïve-EVs) contained two particle populations with peak intensities, 106 nm and 531 nm, but EVs from PLY-challenged cells (PLY-EVs) showed a predominant single population at 220 nm (**Fig. 1C, D**). Zeta potential measurements suggested that the naïve EVs and PLY(0.5)EVs were both negatively charged particles with moderate colloidal stability with values of -16.6 ±2.46 mV and -21.03±3.27 mV respectively.

**Figure 1.**
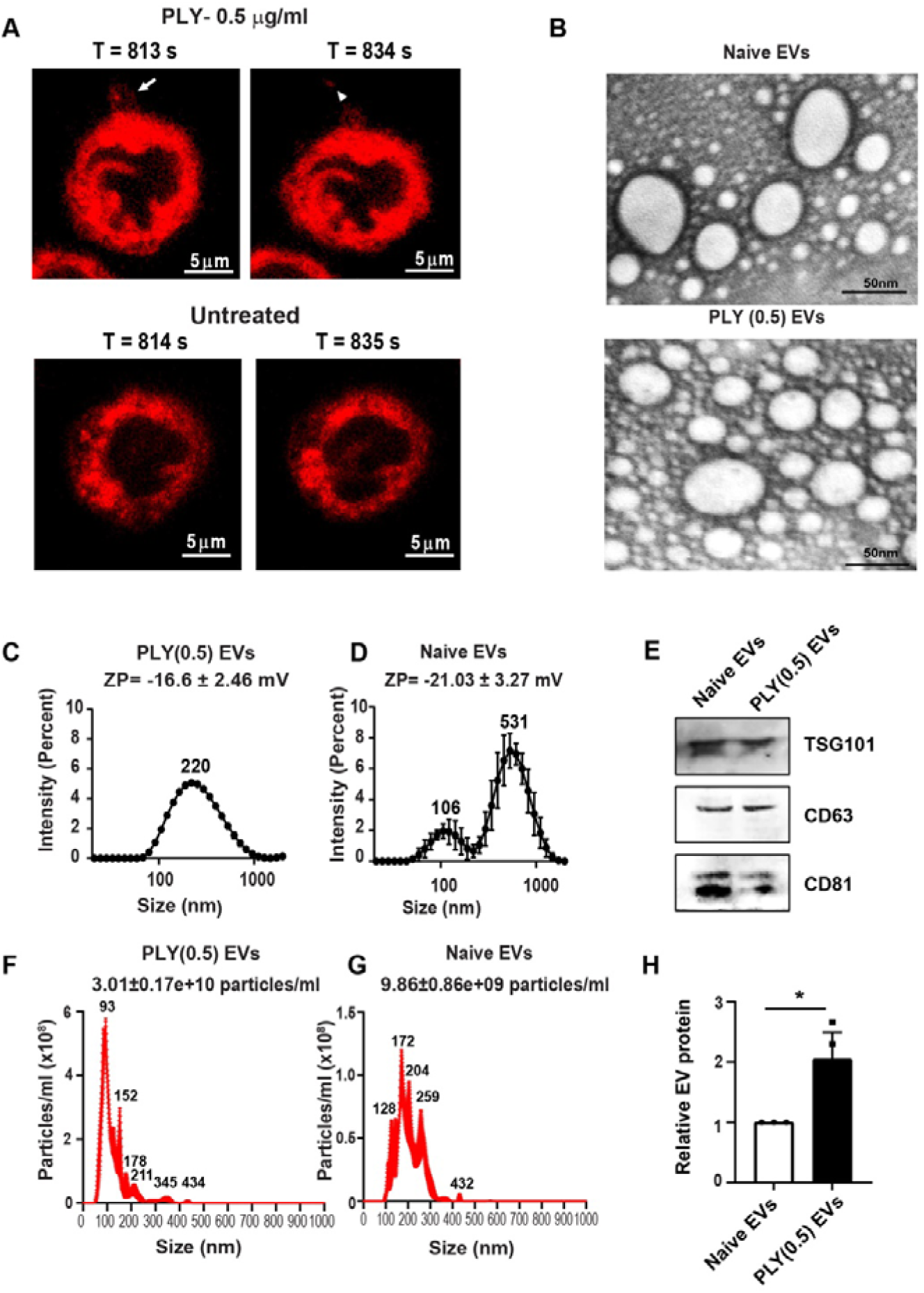
PLY induces EV shedding from human THP-1 monocytes at sublytic doses. **(A)**. Selected time frames from live imaging of Nile-red labelled human THP-1 monocytes stimulated with recombinant PLY (0.5 µg/ml). Arrow indicates membrane protrusions (T=813s) and arrowhead points to the subsequent EV release (T=834s). Scale bars, 5 µm. **(B)**. Representative transmission electron micrographs (N=3) of EVs isolated from PLY-challenged (PLY-EVs) and untreated cells (Naïve EVs). Scale bars, 50 nm. (**C,D**). Dynamic light scattering analysis (N=3) of the particle size distribution showing a single peak of 220 nm for PLY (0.5) EVs and two peaks at 106 nm and 531 nm for naïve EVs. Zeta potential (ZP) values are indicated implying negative surface charge and moderate colloidal stability. **(E)**. Immunoblotting analysis (N=2) showing the presence of characteristic EV markers, TSG101, CD63 and CD81. (**F,G**). NTA analysis (N=5) showing the concentration and size distribution of naïve and PLY(0.5) EVs. (**H**). Quantification of relative total protein content of naïve EVs and PLY(0.5) EVs (N=3) by BCA protein assay. *p<0.05 by t unpaired test.

Next, to confirm the identity of the isolated EVs, we probed the presence of characteristic EV markers by immunoblotting. The vesicles showed the presence of the endosomal sorting complex required for transport (ESCRT-1) subunit, TSG101 and the tetraspanins, CD63 and CD81 (**Fig. 1E**). To quantify the EVs, we performed nanoparticle tracking analysis (NTA) and found that the PLY(0.5)EVs had ∼3-fold higher particle concentration as compared to naïve EVs, suggesting upregulation of vesicle shedding upon PLY challenge (**Fig. 1F, G**). In agreement with the NTA results, BCA quantification of the total EV protein content revealed PLY-EVs had ∼2-fold higher protein abundance relative to naïve EVs (**Fig. 1H**). Henceforth, the naïve EVs and PLY-EVs were normalized with respect to total protein content for all the experiments in the study.

### Transfer of EV-bound PLY to recipient cells upon membrane fusion induces hemolysis and cytotoxicity

To investigate whether cells expel membrane bound PLY into the shed EVs, we performed immunoblotting of EVs as well as whole cell lysates of PLY-challenged cells using monoclonal PLY antibody. We detected PLY on the EVs as well as lysates from PLY-challenged cells, but not on the naïve EVs or untreated cells confirming the specificity (**Fig. 2A**). To examine the localization of PLY on the EVs, we performed immunogold labeling TEM analysis of PLY-EVs and found that PLY was localized on the membrane of PLY-EVs, but not naïve EVs (**Fig. 2B**). Using high-speed confocal microscopy, we also demonstrated the release of GFP-tagged PLY from the membrane of challenged Dil-labelled THP-1 monocytes into the shed EVs (**Video S3**), but not untreated cells (**Video S4)**. Our results are in agreement with the study by Wolfmeier et al., 2016 (*12*), showing the active release of PLY prepores and pores from the damaged cell membrane of PLY-challenged cells as microvesicles. Nevertheless, the fate and downstream consequences of the shed EV-associated PLY have not been investigated. We hypothesized that the EV-membrane localized PLY might be transferred onto the recipient cells upon EV fusion. It has been proposed that vesicle fusion with target cells could result in the insertion of the EV membrane on the target plasma membrane and delivery of EV cargo into the cytoplasm of recipient cells (*23*). To test this hypothesis, we incubated THP-1-monocyte derived DCs with purified PLY- and -naïve EVs for 30 min, washed and subsequently probed the DC membrane lysates for PLY by immunoblotting. Results showed that DCs co-incubated with PLY-EVs, but not naïve EVs showed the presence of PLY, suggesting transfer of PLY from EVs to recipient DCs upon vesicle fusion (**Fig. 2C**). Cells treated with recombinant PLY (rPLY) was used as a positive control. To unequivocally demonstrate the transfer of EV-associated PLY to recipient cells, we purified EVs from THP-1 monocytes challenged with CF647-labelled PLY and stained them with Nile-red. The labelled EVs were subsequently incubated with DCs for 1h. Clearly, colocalization of Nile-red and CF647-PLY in DCs confirmed the specific transfer of EV-bound PLY to recipient DCs upon fusion (**Fig. 2D**). Consistent with the transfer of EV-associated PLY to recipient DCs, we found that PLY-EVs induced higher cytotoxicity in a dose-dependent manner in recipient DCs as compared to naïve EVs (**Fig. 2E**). In order to test whether the PLY-EVs are hemolytic, we incubated normalized protein equivalents of PLY-EVs and naïve EVs with whole blood collected from healthy donors. Results showed that PLY-EVs, but not naïve EVs, induced hemolysis in a dose dependent manner (**Fig. 2F**). rPLY was used as a positive control. To rule out carryover of non-EV bound PLY from culture supernatant into the EV preparations, rPLY containing medium was passed through the 100 kDa ultrafiltration columns akin to EV isolation. The retentate fraction was checked for PLY-hemolytic activity and found to be negative. This confirmed that the measured hemolysis was solely mediated by the EV-bound PLY. Together, our results suggest that PLY-EVs transfer PLY pores to recipient cells upon EV fusion, thereby mediating cytolysis.

**Figure 2.**
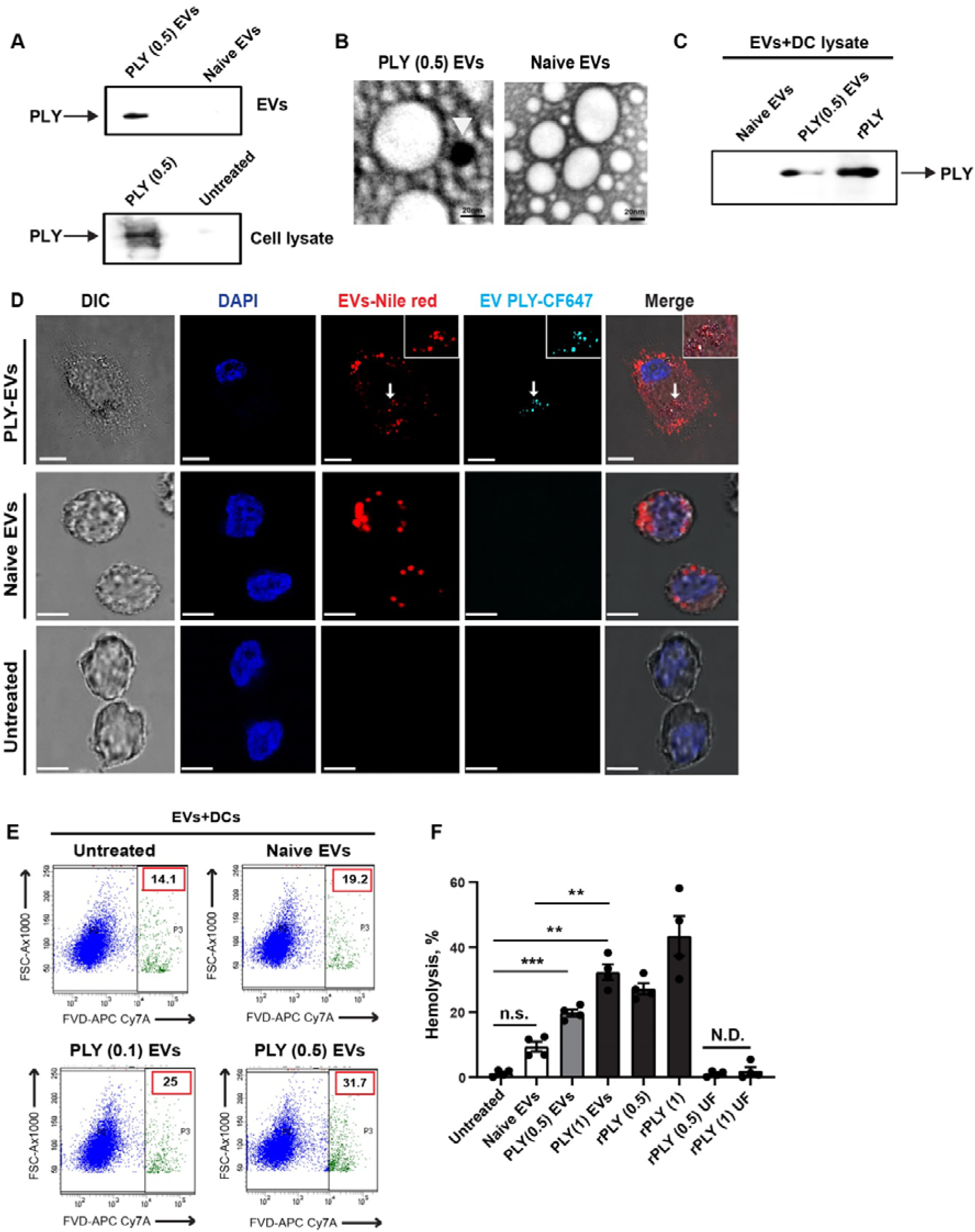
EV-bound PLY induces hemolysis and cytotoxicity upon fusion with recipient cells. **(A)**.Immunoblots of whole cell lysates and EVs isolated from THP-1 cells challenged with PLY (0.5 µg/ml) and untreated cells probed using anti-PLY. (**B**). Immunogold labelling of EVs (N=3) with 12 nm gold nanoparticle conjugated antibody showing the surface localization of PLY on PLY(0.5) EVs, but not naïve EVs (arrow head, 12 nm). Scale bars, 20 nm. (**C**). Immunoblot of DC membrane lysates showing the transfer of PLY upon co-incubation with PLY(0.5)-EVs, but not naïve EVs. Blots are representative of 2 independent experiments. (**D**). Confocal microscopy of DCs incubated with PLY(0.5)EVs for 60 min showing the colocalization of EV-associated CF-647 labelled PLY (cyan) with the EV membrane stain, Nile-red on recipient DCs (magnified in inset). Scale bars, 10 μm. (**E**). Viability staining of DCs treated with PLY EVs (0.1, 0.5) or naïve EVs with fixable viability dye eFluor 780, showing the cytotoxicity of PLY-EVs. Percentage of dead cells are indicated. **(F)**. Hemolysis assay (N=4) showing the dose-dependent induction of red cell lysis upon addition of PLY EVs (0.5, 1) as compared to naïve EVs. Recombinant PLY (rPLY) was used as positive control. rPLY spiked medium passed through 100 kDa ultrafiltration spin columns (rPLY UF) was used as negative control to test for protein carry over into EV preparations. ** represents p< 0.005 and *** represents p<0.0005 by one-way ANOVA with Tukey’s multiple comparisons test. N.D. denotes not detectable.

### PLY-EVs are enriched for proteins involved in immune response and inflammation

To study the differential protein cargo of the PLY-EVs relative to naïve EVs, we performed mass-spectrometric analysis of the EV lysates upon normalization of samples with respect to total protein content. We identified 157 host proteins common to both naïve and PLY EVs, and 38 unique proteins in PLY-EVs (**Fig. 3A**). STRING PPI protein-interaction network analysis revealed that the interactome of PLY-EVs consisted of 54 interactions (edges) linking 36 proteins (nodes) (**Fig. 3B**). The average local clustering coefficient was 0.514 and the protein-protein interaction enrichment p value was 3.12E-08, which demonstrated significant interactions among proteins. We found cytoskeletal proteins-actin and tubulin, RhoA activator, SGK223, membrane ruffling protein, FSCN1 and ESCRT-III protein, CHMP4A that are involved in exosome biogenesis and release through membrane projections. Besides, we found many other proteins involved in cellular stress responses such as heat shock protein, HSPA1B, DNA damage and repair proteins, endonuclease FEN1, p53 regulator, RRP12 and the p38/MAPK14 stress-activated MAPK cascade regulator, TAOK3. The presence of DNA repair proteins in PLY-EVs is in agreement with a previous study by Rai et al., 2016 (*24*), showing that PLY induces DNA double strand breaks in host cells.

**Figure 3.**
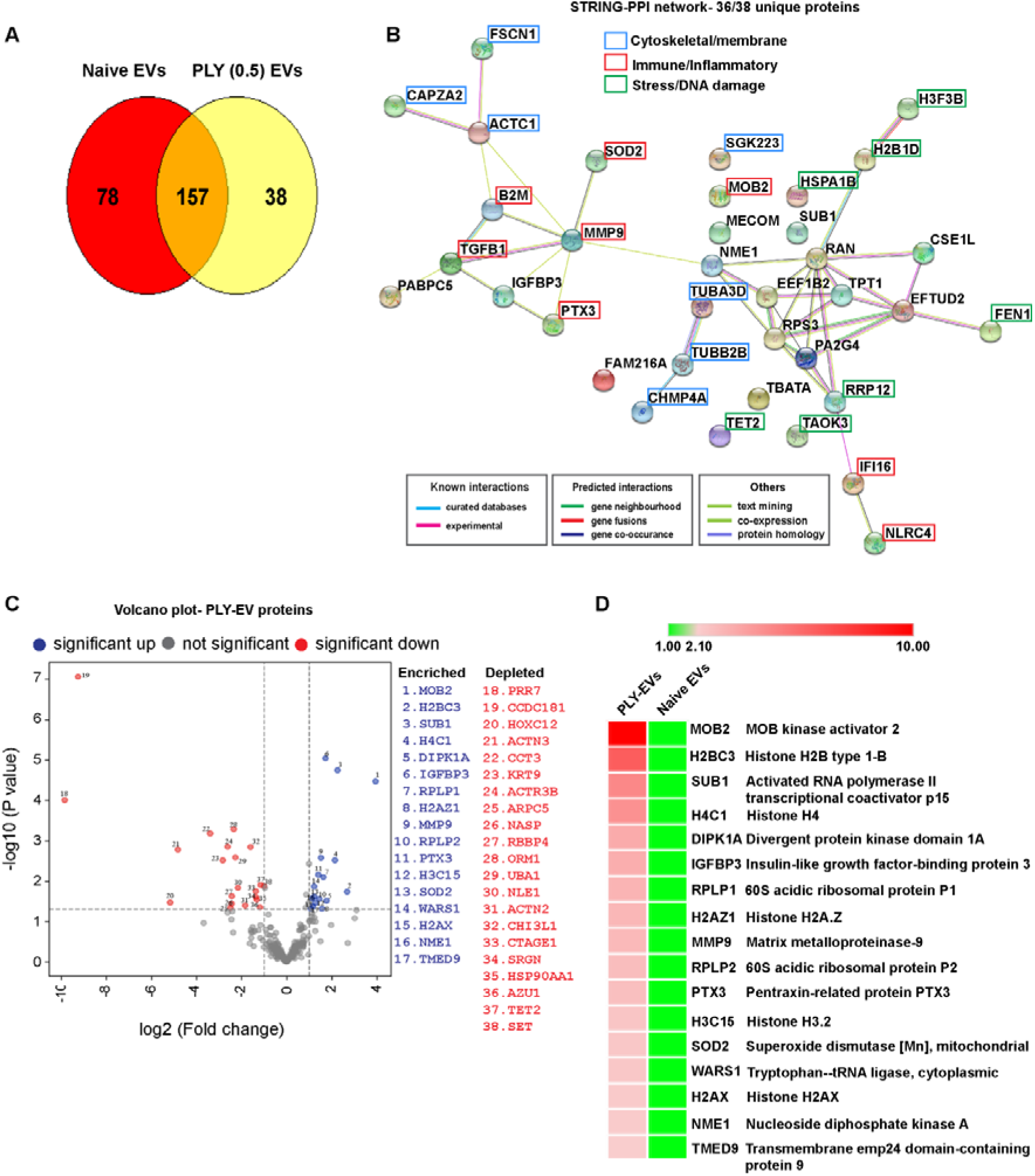
Proteomic analysis of PLY-EV cargo reveals enrichment of immune defense and inflammatory proteins. **(A)**. Venn-diagram plot of proteins identified in naïve and PLY EVs by LC-MS/MS analysis. (**B**). STRING protein interaction network analysis of proteins unique to PLY-EVs. A total of 54 interactions (edges) linking 36 proteins (nodes) were observed. The average local clustering coefficient was 0.514 and protein-protein interaction enrichment p value was 3.12E-08 which demonstrated significant interactions among proteins. (**C**). Volcano plot analysis of differential protein abundance in PLY (0.5) EVs. Blue and red shows differentially enriched and depleted proteins respectively with fold change > 2 and p < 0.05. (**D**). Heat map showing the significantly enriched proteins in PLY (0.5) EVs compared to naïve EVs. Proteomics data are representative of 3 biological replicates.

Differential protein content of proteins in PLY-EVs relative to naïve EVs was classified based on the cut off for fold change> 2 and p < 0.05 to create the volcano plot (**Fig. 3C**). Importantly, we identified several proteins implicated in immune defense and inflammatory signaling exclusively in PLY-EVs, such as the gamma interferon inducible protein, IFI16, MHC-I component B2M, NLR family CARD domain containing protein 4 (NLRC4), complement activating long pentraxin, PTX3, matrix metalloproteinase, MMP9 and superoxide dismutase, SOD2. Heat map analysis of the significantly enriched proteins identified the serine threonine kinases activator, MOB2 that regulates DNA damage response (*25*) and cell cycle progression (*26*) in cancer cells (**Fig. 3D**). MOB2 has been shown to activate the kinases, NDR1/2 that play pivotal roles in inflammatory cytokine signaling and antimicrobial innate immune response (*27*). Besides, we identified canonical and histone variants such as H2B, H4, H3.2, H2A.Z and H2AX to be significantly enriched in PLY-EVs as compared to naïve EVs. Histones have been reported to be enriched in exosomes (*28*), consistent with the theory that cells get rid of damaged DNA by packaging them into exosomes to prevent type I IFN release via the c-GAS STING pathway (*29*). Besides their well-established roles in nuclear chromatin organization, cationic histones have been proposed to have antimicrobial activity by disrupting bacterial membrane (*30*). Hence, the consequence of histones that are enriched in the PLY-EVs, on the bacterial fitness during an infection remains to be investigated. Taken together, our results identified that PLY-EVs contain antimicrobial, stress response and inflammatory proteins that could modulate the host response to infection.

### PLY-EVs induce dendritic cell maturation and inflammatory cytokine release upon internalization

To investigate the effect of PLY-EVs on immune cells, we tested their internalization by DCs, which are the major antigen presenting cells and the resulting consequences on DC activation. We incubated CFSE-labelled PLY-EVs and naïve EVs with human monocyte-derived DCs for 24h and visualized their internalization by confocal microscopy. Results showed that PLY-EVs were internalized to a greater extent by the DCs than naïve EVs (**Fig. 4A**). The EV uptake was quantified by flow cytometry, confirming the higher uptake of PLY-EVs relative to naïve EVs and in a dose-dependent manner (**Fig. 4B, C**). Similar to PLY-EVs, EVs derived from THP-1 monocytes infected with PLY-expressing *S. pneumoniae* serotype 4 strain, T4R (T4R-EVs), showed higher internalization by A549 human alveolar basal epithelial cells as compared to EVs from cells infected with isogenic PLY mutant strain, T4RΔPLY (T4RΔPLY EVs) (**Fig. S3A, B**). Interestingly, we also found that PLY-EVs, but not naïve EVs induced dendritic processes that are characteristic of maturation in stimulated immature DCs (**Fig. 4D**). To verify the DC maturation induced by PLY-EVs, we quantified the expression of typical DC activation markers by flow cytometry. Consistent with Fig. 4D, we found that DCs incubated with PLY-EVs, but not naïve EVs upregulated the surface expression of co-stimulatory molecules, CD80 and CD86, as well as the maturation marker, CD83 (**Fig. 4E-G**).

**Figure 4.**
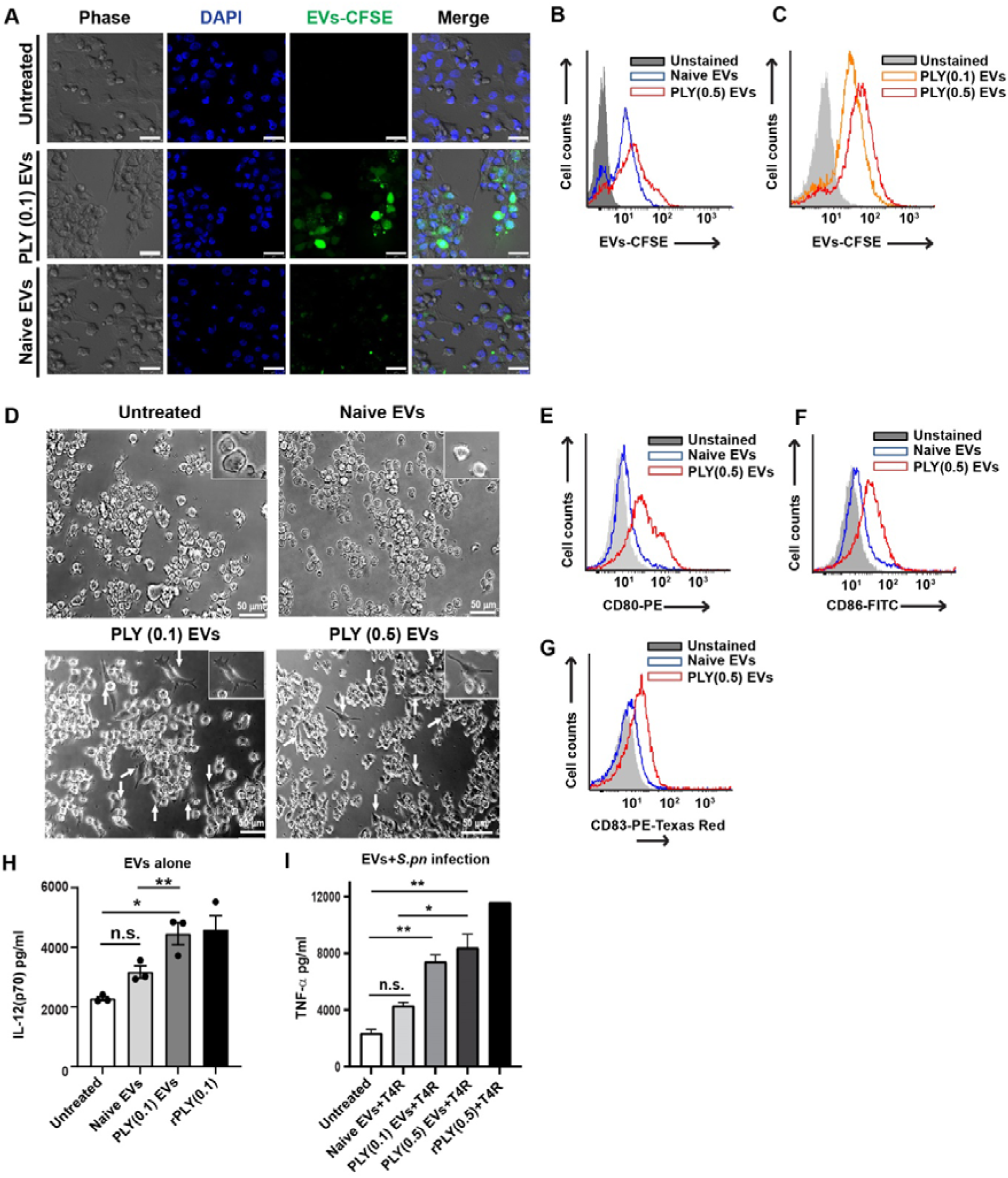
PLY-EVs are internalized by DCs and induce maturation and inflammatory cytokine response. **(A)**. Confocal microscopy showing the internalization of CFSE-labelled PLY (0.1) and naïve EVs (green) by DCs at 24h post treatment. Scale bars, 25 µm. (**B**). Flow cytometry histograms (N=3) to quantify the DC uptake of CFSE labelled PLY (0.5) EVs and naïve EVs. (**C**). Dose-dependent uptake of PLY (0.1, 0.5) EVs by DCs. (**D**). Phase-contrast microscopy of immature DCs treated with PLY (0.1, 0.5) EVs and naïve EVs. Arrows indicate matured DCs (magnified in inset). Scale bars, 50 μm. (**E-G**). Flow cytometry histograms (N=3) to quantify the expression of (**E**) CD80, (**F**) CD86 and (**G**) CD83 on DCs treated with PLY (0.5)- and naïve-EVs. (**H, I**). Cytokine ELISA showing the levels of secreted (**H**) IL-12(p70) from DCs treated with PLY (0.1) EVs or naïve EVs alone and (**I**) TNF-α from DCs pre-treated with PLY (0.1,0.5) or naïve-EVs for 24h followed by subsequent infection with *S. pneumoniae*, T4R strain. Recombinant PLY (0.5 µg/ml) was used as positive control. Data are representative of 2 independent experiments. * indicates p<0.05 and ** p<0.005 by one-way ANOVA with Tukey’s multiple comparisons test. n.s. denotes not significant.

Next, we tested the consequence of EV internalization by DCs on the latter’s inflammatory cytokine response. We found that PLY-EVs treated DCs produced higher levels of the inflammatory Th1 cytokines, IL-12 (**Fig. 4H**) and TNF-α (**Fig. S3C**), as compared to cells treated with naïve EVs. Based on our findings showing immunostimulatory potential of PLY-EVs, we hypothesized that EVs shed from PLY-challenged cells could prime bystander naïve cells to stronger response against infection. To test this, DCs were pre-treated with PLY-EVs or naïve EVs for 24h and subsequently infected with *S. pneumoniae*, T4R strain, following which the secretion of the pro-inflammatory cytokines, IL-12 and TNF-α were measured. Results showed that DCs primed with PLY-EVs produced significantly higher amounts of TNF-α (**Fig. 4I**) and IL-12 (**Fig. S3D**) upon infection, when compared to naïve-EV treated cells. In agreement, we found that EVs derived from THP-1 monocytes infected with PLY-expressing T4R strain, induced higher TNF-α from DCs as compared to EVs from T4RΔPLY infected cells (**Fig. S3E**). Taken together, our findings show that PLY-EVs activate DCs upon internalization and alert the cells to infection.

### PLY-EVs induce inflammation and toxicity *in vivo*

To study the toxicity and pathophysiological effects of the toxin-induced host EVs *in vivo*, we first employed a zebrafish (*Danio rerio*) model in which CFSE-labelled naïve- and PLY(0.5)EVs were microinjected into the Duct of Cuvier of day 3 post-fertilization embryos. Zebrafish is a widely accepted model system for human infectious diseases, due to their optical transparency and similarity with the human immune system (*31*) and have been previously validated by us (*32*) and others (*33*) as a suitable model to study pneumococcal pathogenesis. We found that zebrafish administered with PLY-EVs, but not naïve EVs developed severe pericardial edema at day 1 post injection confirming the toxicity of PLY-containing EVs (**Fig. 5A**). PBS treated embryos served as the mock injected group. The PLY-EV administered group also showed a progressive decline in the heartbeat rate as compared to naïve EV and mock-injected control groups, indicative of cardiac dysfunction (**Fig. 5B**). In conjunction with the severe cardiac dysfunction and edema formation, the PLY-EV treated zebrafish also showed a significantly lower survival compared to naïve EV and mock-injected group, with only 30% of the embryos surviving at day 4 post injection (**Fig. 5C**). Moreover, zebrafish administered with PLY-EVs showed ∼2.5 log fold higher expression of the pro-inflammatory cytokine, TNF-α relative to naïve EV treated group (**Fig. 5D**).

**Figure 5.**
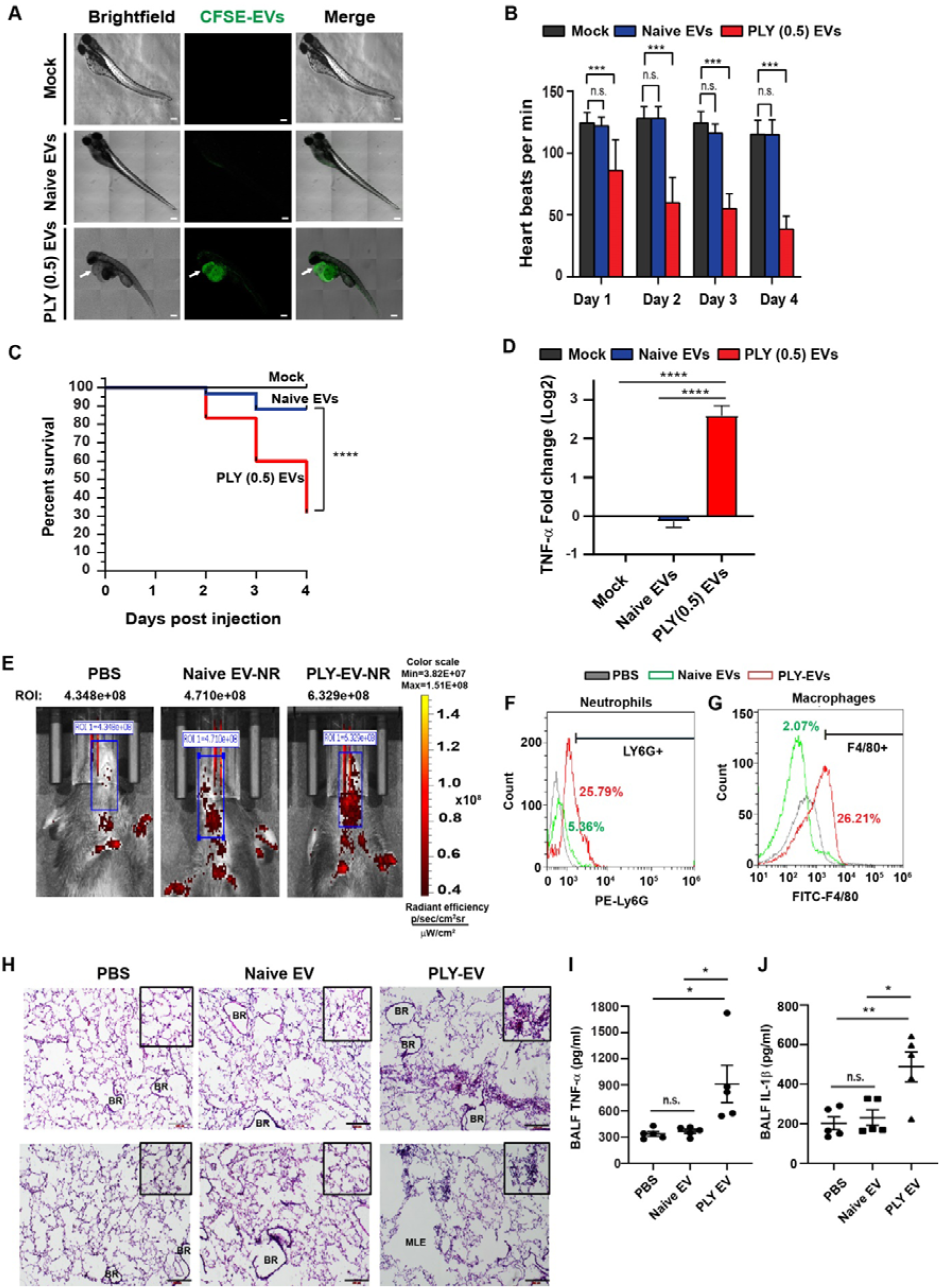
PLY-EVs induce inflammation and toxicity *in vivo*. **(A)**. Confocal microscopy of zebrafish embryos (N=60) injected with CFSE labelled naïve or PLY(0.5)EVs (green) at day 1 post injection showing the development of pericardial edema in the PLY-EV treated group (arrows). Scale bars, 100 μm. (**B**). Heartbeat and (**C**) survival graph of zebrafish (N=60) injected with naïve or PLY (0.5)EVs was monitored upto 4 days post injection. *** in panel B denotes P< 0.01 by Mann-Whitney (two-tailed) test. **** in panel C denotes P<0.0001 by Log-rank (Mantel-Cox) test. n.s. denotes not-significant. (**D**). Real-time PCR experiment (N=3) showing the relative expression of TNF-α mRNA in zebrafish treated with naïve or PLY(0.5) EVs at day 4 relative to mock injected group. **** denotes P<0.0001 by one-way ANOVA with Tukey’s multiple comparisons test. (**E**). IVIS imaging of C57BL/6 mice adoptively transferred with Nile-red labelled naïve EVs or PLY-EVs. PBS treated mice served as negative control. ROI intensity values are indicated showing higher intensity of PLY-EVs in the respiratory tract. (**F, G**). Flow cytometry analysis of neutrophils (Ly6G^+^) and inflammatory macrophages (F4/80^+^) in BALF of mice (n=6) adoptively transferred with naïve EVs or PLY-EVs at 24 h. (**H**). Hematoxylin and eosin (H&E) staining of mouse lungs (n=6) at 24h post adoptive transfer of PLY-EVs, naïve EVs or PBS. Mice adoptively transferred with PLY-EVs showed tissue microlesions (MLE) and immune infiltration in the alveolar spaces indicative of PLY-induced tissue damage (magnified in the inset). BR-bronchiole; MLE- microlesions. Scale bars, 200 µm. (**I-J**) Levels of pro-inflammatory cytokines (**I**) TNF-α and (**J**) IL-1β in the BALF of mice (n= 5 mice/group) adoptively transferred with naïve EVs or PLY-EVs were measured post-sacrifice at 24h. PBS treated mice served as negative control. Mouse BAL flow cytometry and histology data are representative of 3 independent experiments. * in panels I, J indicates p<0.05 and ** p<0.005 by one-way ANOVA with Tukey’s multiple comparisons test. n.s. denotes not significant.

Next, we used a mouse model to investigate the toxicity of PLY-induced EVs *in vivo*. To mimic PLY-induced acute lung injury during infection, 6-8 weeks old C57BL/6 mice were intranasally instilled with 0.25 μg of purified recombinant PLY in sterile PBS. The PLY dose was chosen based on a previous study showing the induction of lung hyperpermeability and acute lung injury in mice upon aerosol administration of 0.25 µg of PLY (*34*). Zafar et al., showed that administration of recombinant PLY dose of 200 ng/day induced shedding of pneumococci and mucosal inflammation (*35*). Intranasal administration of PLY at 10 ng/g body weight (corresponding to 0.25 μg for 6-8 week mice) restored virulence of an avirulent ST306 pneumococcal clone expressing a non-cytolytic PLY variant (*36*). Based on the above studies, we tested two intranasal doses of 0.25 and 0.5 μg PLY, but we observed mortality at 0.5 μg dose. Hence, we used intranasal dose of 0.25 µg PLY in 50 μl PBS to model PLY-induced acute lung injury. We found that by 18h post PLY administration, mice showed a significant increase in lung vascular permeability as confirmed by extravasation of the intravenously administered plasma tracer dye, Evans blue through IVIS imaging (**Fig. S4A**). This was concomitant with the presence of GFP-tagged PLY in the lungs of treated mice (**Fig. S4B**). PLY-induced lung inflammation was also confirmed by higher infiltration of neutrophils and macrophages into the lungs (**Fig. S4C**). Histological staining revealed regions of tissue microlesions and immune cell infiltration into the alveolar spaces of the PLY-treated mice (**Fig. S4D**). At 18h, EVs were isolated from murine bronchoalveolar lavage fluid (BALF) collected by lung perfusion post mortem. The BALF EVs isolated from lungs of untreated and PLY-treated donor mice were administered intranasally to healthy recipient mice upon normalizing the total protein concentration of EVs. At 24h post EV transfer, the distribution of the Nile-red labelled EVs in the murine respiratory tract was visualized by IVIS imaging. Results showed that the administered PLY-EVs showed higher signal intensity even at 24h as compared to naive-EVs derived from untreated mice (**Fig. 5E**). PBS administered mice served as negative control. Flow cytometry analysis of the murine BALF revealed an enrichment of neutrophils (**Fig. 5F**) and macrophages (**Fig. 5G**) in lungs of mice treated with PLY-EVs, in comparison to naive-EV and PBS treated groups. This was also confirmed by H&E lung histology staining showing microlesions and inflammatory lymphocyte infiltration into the alveolar spaces of PLY-EV treated mice (**Fig. 5H**). Naive-EV and PBS treated mice had normal lung morphology with intact alveolar space and absence of inflammatory cells. Further, PLY-EV treated mice showed significantly higher levels of the pro-inflammatory cytokines, TNF-α (**Fig. 5I**) and IL-1β (**Fig. 5J**) in the BALF at 24h when compared to naive-EV and PBS treated mice. Taken together, our results show that the PLY-EVs induce toxicity and inflammation *in vivo*.

## Discussion

Pneumolysin (PLY), the β-barrel pore-forming toxin released by the respiratory tract pathogen, *S. pneumoniae* binds to membrane cholesterol of eukaryotic host cells and oligomerizes to form transmembrane pores, resulting in lytic cell death (*10*). In this study, we found that stimulation of human monocytes with sublytic doses of recombinant PLY upregulates extracellular vesicle shedding from the challenged cells. Further, we also showed that the shed EVs harbour membrane-localized PLY, which is in agreement with previous studies that showed the expulsion of PLY pores from cells though microvesicle shedding (*12, 14, 37*). However, the downstream effects of the shed PLY-EVs on the host are unknown and it is believed that the pore-forming toxins are completely neutralized upon capture by synthetic cholesterol-containing liposomes (*38-40*). However, neither of these studies investigated the downstream effects of the toxin-soaked liposomes. Moreover, synthetic liposomes differ from host cell-derived EVs with regards to membrane lipid composition as well as the presence of surface proteins that mediate cell tropism and fusion with target cells (*41*). Our findings reveal that EVs shed from PLY-challenged cells, harbour membrane bound PLY that induces dose-dependent hemolysis and cytotoxicity in recipient cells upon fusion. This was confirmed by experiments showing the incorporation of labelled EV-membrane associated PLY into recipient DCs upon co-incubation. Our findings are in agreement with studies showing the transfer of EV-bound MHC-peptide complex to antigen-presenting cells upon membrane fusion in a process termed as cross-dressing (*23, 42*).

Proteomic analysis of the PLY-EV cargo revealed the presence of key proteins involved in innate immune response and inflammation such as the gamma interferon inducible protein, IFI16, long pentraxin related PTX3, the Nod-like receptor, NLRC4, matrix metalloproteinase, MMP9 and superoxide dismutase, SOD2. The long pentraxin, PTX3 is a complement regulating protein that was reported to be upregulated upon pneumococcal infection (*43*). Besides, we also found the presence of the matrix metalloproteinase, MMP-9 specifically in PLY-EVs. PLY has been reported to promote MMP release from neutrophils and plays an important role in the progression of severe pneumococcal disease (*44*). PLY triggers oxidative stress (*45*), hence it is conceivable that EVs contain superoxide dismutase to protect recipient cells from oxidative damage. Through oxidative damage, PLY also induces DNA double strand breaks, and results in the noncanonical histone H2 phosphorylation at the Ser-129 residue (H2AX). In agreement, we found that PLY-EVs contain the modified histone, H2AX suggesting that the cells get rid of damaged DNA in the cytoplasm through vesicles to maintain cellular homeostasis, consistent with Takahashi et al., 2017 (*29*). Besides, we identified several other histones, H2B, H3 and H4 to be significantly enriched in PLY-EVs, relative to naïve EVs. Recently, mammalian histones have shown to exhibit synergy with antimicrobial peptides such as LL-37, by affecting bacterial membrane stability and inhibiting transcription (*46*), suggesting that the PLY-induced EVs could be bactericidal. Taken together, our results suggest that EVs released from host cells harbour potential damage associated molecular pattern molecules which may serve to alert the host of infection and mount an anti-microbial and inflammatory response. Further studies are necessary to determine the effects of these EVs on bacterial clearance by the host immune system.

Upon fusion with target cells, EVs transfer the cargo into the recipient cells. We found that PLY-EVs were better internalized by DCs as compared to naïve EVs. This could be explained by the fact that PLY induced endocytosis via MRC-1 receptor (*47*) in DCs and dynamin-dependent endocytosis in glial cells (*15*). Upon internalization, the PLY-EVs induced DC maturation, consistent with the upregulation of co-stimulatory molecules and maturation markers. Naïve EVs shed from untreated cells were also internalized, but did not induce DC activation supporting the notion that exosome-derived self-peptides maintain peripheral tolerance (*48*). Importantly, pre-treatment of DCs with PLY-EVs, elicited higher inflammatory cytokine response upon infection with *S. pneumonia*e, compared to naïve EVs. This suggests that the EV-associated PLY activates bystander cells for a robust response against infection. We also verified the immunostimulatory effect of PLY-EVs with EVs from cells infected with PLY-expressing *S. pneumoniae*, T4R strain as well as the isogenic PLY mutant strain, T4RΔply. Our data prompts us to speculate that EVs shed in response to bacterial toxin may elicit protection against infection due to their ability to prime DC responses. It also remains to be studied whether the EVs may harbour MHC-bound antigens to T cells leading to antigen specific T cell responses.

Using two *in vivo* models, zebrafish and mice, we demonstrated that PLY-EVs induce toxicity and inflammation. Zebrafish embryos administered with PLY-EVs showed severe pericardial edema, impaired heart function and reduced overall survival compared to naïve EVs. Circulating PLY has been shown to induce cardiac injury during pneumococcal infection(*49, 50*). Our study identifies a previously unrecognized mechanism for exacerbation of toxicity through EVs shed from PLY-challenged host cells. We developed a PLY-induced acute lung injury mice model and showed that EVs isolated from lungs of these mice induced severe inflammation when transferred to healthy recipients. Overall, our study proposes a mechanism by which host vesicles shedded in response to bacterial toxins drive toxicity and inflammation (**Fig. 6**). Our findings shed light on the role of host-derived vesicles during pneumococcal pathogenesis and their potential applications in EV-based diagnostics as well as therapeutics.

**Figure 6.**
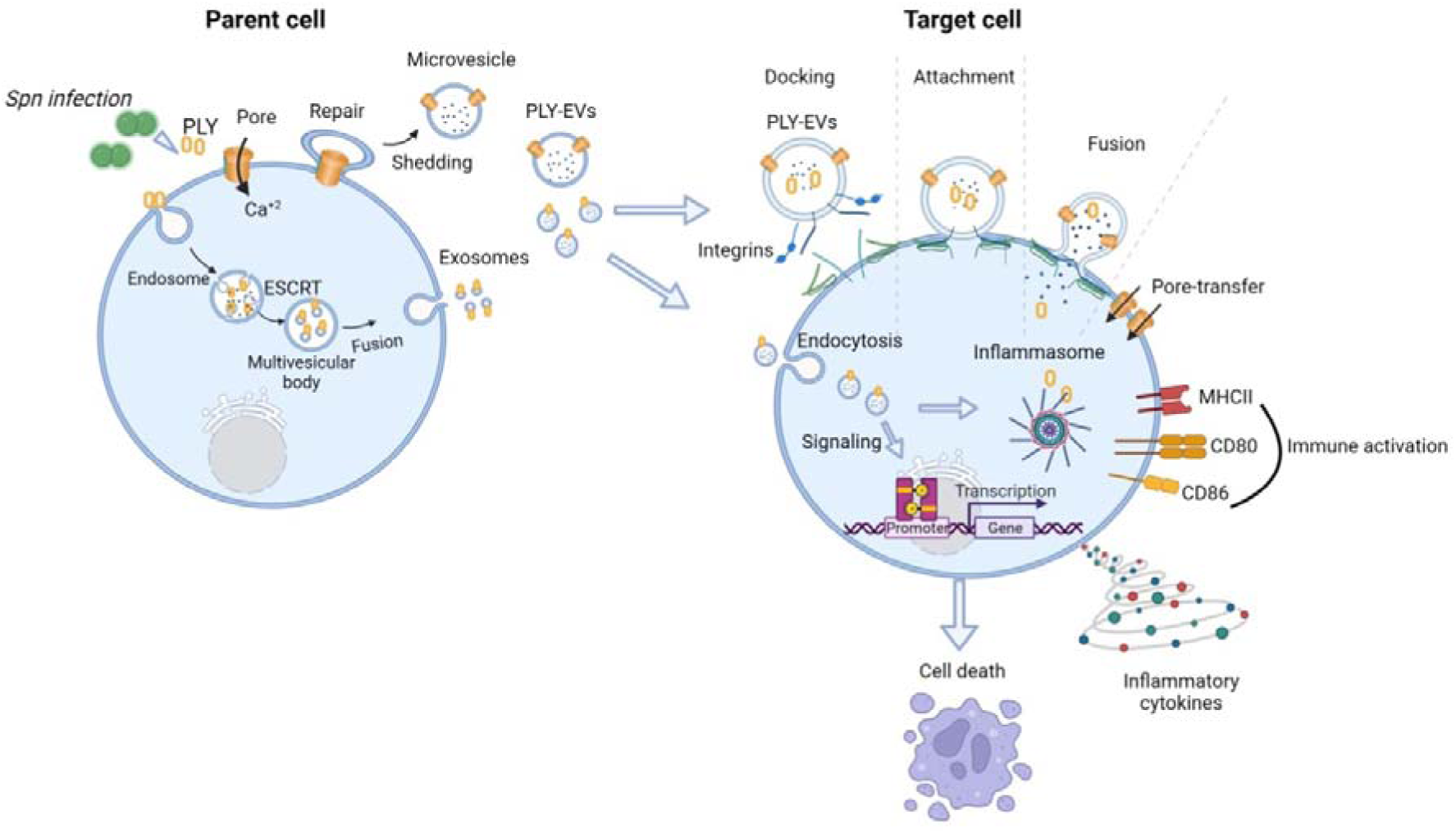
Extracellular vesicles shedded in response to pneumolysin induce toxicity and inflammation in the host. During infection, *S. pneumoniae* releases the pore-forming toxin, PLY that oligomerizes to form pores on the plasma membrane of eukaryotic host cells. At sublytic toxin doses, Ca^2+^-dependent intrinsic cell membrane repair mechanisms are proposed to remove toxin pores by shedding the damaged membrane as microvesicles. Concurrently, the PLY can also be endocytosed into endosomes that form multivesicular bodies through internal budding mediated by ESCRT protein complex and fuse with the plasma membrane to release exosomes bearing PLY. The toxin-bearing extracellular vesicles (EVs) contain docking receptors such as integrins that enable fusion with target cells. Here, we show that EVs shed from PLY challenged host cells harbour membrane bound PLY and upon fusion with target cells, the EV membrane gets integrated into the recipient cells, thereby transferring the PLY-pores and inducing cytotoxicity. We also identified the unique protein cargo of PLY-EVs that could exert inflammatory effects in the host. Further, we show upregulation of costimulatory molecules and pro-inflammatory cytokines on dendritic cells exposed to PLY-EVs. We verify the pathophysiological significance of toxin-induced EVs *in vivo* using two models, zebrafish and mouse.

## Methods

### Cell cultures

Human THP-1 monocytes (ATCC TIB-202) were cultured in RPMI 1640 medium (HiMedia) supplemented with 10% fetal bovine serum (FBS) (Invitrogen). Human alveolar basal epithelial cells, A549 (ATCC CCL-185) was grown in DMEM (HiMedia) supplemented with 10% FBS. Cell cultures were maintained in 5% CO_2_ at 37°C. The cell lines were procured from the national cell repository at the National Centre for Cell Science, Pune and were authenticated using short tandem repeat analysis and mycoplasma testing. Monocyte-derived DCs were differentiated by stimulating THP-1 cells with human GMCSF (1500 IU/ml) and IL-4 (1500 IU/ml) (Peprotech) for 5 days with half the medium replaced at day 3. To induce maturation, TNF-α (2000IU/ml, Peprotech) and Ionomycin (200 ng/ml) was added at day 5 and further incubated for 2 days. The DCs were verified for surface expression of characteristics markers, CD80, CD86 and CD83 by flow cytometry.

### Bacterial cultures and infection

*Streptococcus pneumoniae* serotype 4 unencapsulated strain, T4R and the isogenic pneumolysin mutant, T4RΔPLY (a kind gift from Prof. Birgitta Henriques Normark, Karolinska Institutet, Stockholm through MTA) were grown overnight on soyabean casein digest agar plates with 5% sheep blood (HiMedia) in 5% CO_2_ at 37°C. For infection, liquid cultures were grown in brain heart infusion broth (HiMedia) at 37°C to OD of 0.4 and pelleted at 5000 rpm for 5 min. All infection experiments were performed with prior approval from the Institutional biosafety committee, RGCB (Ref. 51/IBSC/karthik/202151) and Review Committee on Genetic Manipulation, Department of Biotechnology, Govt. of India (Ref. BT/IBKP/363/2020).

### Isolation of EVs from PLY-challenged and infected cells

To pre-deplete serum-derived exosomes, FBS supplemented RPMI medium was centrifuged at 100,000g overnight and passed through 0.22-μm filter. For isolation of PLY-EVs, 6 x 10^7^ THP-1 monocytes were stimulated with 0.1 μg/ml and 0.5 μg/ml of purified recombinant His-tagged PLY (a kind gift from Prof. Anirban Banerjee, Indian Institute of Technology Bombay) in EV-depleted medium for 24 h at 37□. The percentage of dead cells was quantified by staining with the Fixable Viability Dye eFluor 780 (eBioscience, 1:50,000 dilution) following the manufacturer’s protocol. For MTT cell viability assay, 10-fold serial dilutions from 1 x 10^6^ to 1 x 10^3^ cells/mL were incubated with 0.5 mg/mL MTT dye (Sigma) for 2 h, solubilized using 100 μl DMSO and MTT dye absorbance at 570 nm was measured using Varioskan LUX multimode microplate reader (ThermoFisher). For infection-derived EVs, cells were infected with *S. pneumoniae*, T4R or isogenic PLY mutant strain, T4RΔPLY in EV-depleted medium for 2 h at MOI of 10, following which the extracellular bacteria were removed by PBS wash and gentamicin (0.1 mg/ml) treatment for 1h. At end of time point, culture supernatants were differentially centrifuged at 300g for 10 min, 2000g for 20 min and 10,000g for 30 min to pellet cells, debris, apoptotic bodies and larger microvesicles respectively. The conditioned supernatant was passed through 0.45-μm filter and concentrated using 100 kDa cut-off centrifugal filter units (Amicon). The EVs were precipitated using ExoQuick-TC reagent (System Biosciences) following the manufacturer’s instructions and depleted of serum proteins using 100 kDa spin columns. The EVs were resuspended in PBS and stored at -80°C.

### Live-imaging of EV shedding from cells

2×10^5^ THP-1 cells were labelled with the lipophilic membrane dye, Nile red (40 µg/ml, Invitrogen) and seeded on poly-lysine coated μ-Slide 8 well chambered slides (Ibidi). The cells were stimulated with 0.5 µg/ml PLY or PBS in an enclosed microscope chamber maintained at 37°C and 5% CO_2_. Images were acquired at 5s interval for a total of 20 min under the 63x oil objective of Leica SP8 laser confocal microscope (Leica Microsystems). Videos were processed using the Leica LASX software at 2fps.

### Dynamic light scattering and nanoparticle tracking analysis

Particle size distribution and zeta potential measurements were performed using the Zetasizer particle analyser (Malvern laboratory, Aimil, Bangalore). Triplicate measurements were performed with standard settings (Refractive Index 1.330, Viscosity 0.8872, Temperature 25°C). The NTA analysis was performed using the Nanosight NS300 at Malvern PANalytical Application Lab, Delhi using 20x microscope attached to a CMOS camera. The EVs were diluted in PBS to achieve a minimum of 20 particles per frame and maximum of 120-150 particles per frame.

### Immunogold labelling and transmission electron microscopy (TEM)

For transmission electron microscopy, 10 µl of purified EVs were adsorbed onto Formvar coated copper grids (Electron Microscopy Sciences), blocked with 1% BSA and incubated with mouse anti-PLY (1:15, Abcam) followed by 12 nm gold conjugated goat anti-mouse IgG (Electron Microscopy Sciences) (1:20 dilution). Samples were fixed using 2% glutaraldehyde and contrasted using UranyLess stain (Electron Microscopy Sciences). After drying, the grids were observed under the JEOL 1011 transmission electron microscope at 100 KV.

### Immunoblotting for detection of EV markers and PLY

The EVs were lysed in RIPA lysis buffer and the total protein was quantified using the BCA protein assay kit (Pierce). The lysates were resolved on 12% SDS-PAGE gel and transferred to PVDF membrane. The blots were blocked using 5% BSA for 1h and probed with mouse anti-CD81 (1:500, ab79559), rabbit anti-CD63 (1:1000, ab271286), rabbit anti-TSG101 (1:1000, ab125011) and mouse anti-PLY (1:500, ab71810) overnight at 4°C. HRP-conjugated anti-mouse IgG and anti-rabbit IgG (1:3000, Biorad) were used and imaged using the imaged using Azure 600 gel imaging system (Azure Biosystems).

### Hemolysis assay

Peripheral blood was collected from healthy donors by a trained phlebotomist and the study was approved by the Institutional human ethical committee, RGCB (Ref. IHEC/12/2020_E/14). The EV suspension in PBS was co-incubated with 2% blood suspension in PBS containing 1 mM DTT for 1 h at 37□ in V-shaped bottom 96-well plates (Tarsons). Purified recombinant PLY (0.5 μg/ml and 1 μg/ml) was used as positive control. To control for carry-over of non-EV PLY from culture supernatant into the EV preparations, rPLY spiked culture medium was passed through the 100 kDa ultrafiltration spin columns as performed during EV isolation and the retentate fraction was checked for PLY-hemolytic activity. The plates were centrifuged at 400xg for 15min and the absorbance of supernatant was measured at 540 nm using the Varioskan LUX multimode microplate reader (Thermo Fisher). Triton X-100 (0.2%) was used as 100% lysis control and all the values were normalized to this control and relatively expressed as % hemolysis.

### CFSE labelling of EVs and internalization by DCs

EVs were labelled with CFSE dye (10 μM, Invitrogen) for 1 h at 37°C and the excess dye was removed using 100 kDa MWCO centrifugal spin columns. The CFSE labelled EVs were incubated with THP-1 monocyte-derived DCs grown on coverslips for 24 h at 37°C. The cells were fixed with 4% PFA for 10 min and mounted on glass slide using the ProLong Gold anti-fade mounting medium with DAPI (Invitrogen). Images were acquired under the 63x oil objective of Leica SP8 laser confocal microscope (Leica Microsystems).

### Mass-spectrometry and data analysis

For LC-MS/MS analysis, EVs were isolated as described previously in culture medium lacking FBS to exclude serum proteins. The EVs were lysed by direct boiling in Rapigest SF (Waters) and relative total protein concentrations normalized to ∼1 mg/ml by BCA assay. The samples were reduced using 100 mM 1, 4-Dithiothreitol (Sigma Aldrich), alkylated using 200 mM iodoacetamide (Sigma Aldrich) and digested overnight with MS grade trypsin (Sigma Aldrich) in the ratio 1:25. The injection volume was 2.0 µl and all the samples were injected in duplicate into the nano ACQUITY UPLC chromatographic system (Waters) and MS runs were performed using the Synapt G2 High-Definition MS System (Waters). The data was analyzed using the Progenesis QI for Proteomics v4.2 (Non-Linear Dynamics, Waters). Relative protein quantification was performed using the Hi3 label free method. The naive EVs and PLY(0.5) EVs datasets were analyzed with Exocarta and Vesiclepedia databases to distinguish between classical (endoplasmic reticulum/golgi) and non-classical (such as exosomes and microvesicles). The desired protein lists thus obtained were then computed for their assignment and enrichment levels using different functional categories, including KEGG metabolic pathways and Gene Ontology (GO) terms. A functional protein-protein interaction network of differentially expressed proteins in PLY(0.5) EVs was constructed based on the STRING database using Cytoscape. A Venn diagram showing the distribution of shared and specific proteins among the naive EVs and PLY-EVs was generated using the online Venn diagram program (http://bioinformatics.psb.ugent.be/webtools/Venn/). Volcano plot and heatmap showing the differentially enriched proteins in PLY(0.5) EVs as compared to Naïve EVs were generated using R software. Protein-interaction network analysis of enriched proteins in PLY(0.5) EVs was performed using the STRING functional annotation protein interaction database.

### Flow cytometry of EV uptake and DC maturation

To quantify EV uptake, CFSE-labelled EVs were incubated with 2×10^5^ cells for 24 h and the samples were washed and fixed with 4% PFA. The EV uptake was analyzed on the FITC channel using the BD FACS Aria III cytometer (BD Biosciences). For DC maturation assays, immature day 5 stage DCs were incubated with EVs for 24h and stained using PE-mouse anti-CD80, Clone L307.4 (BD Biosciences), FITC-Mouse anti-CD86 Clone-2331(BD Biosciences), PE-anti-human CD83 clone HB15 (Biolegend) as per manufacturer’s instructions. Cells were stained with fixable viability dye (FVD eFluor 780, eBiosciences) to exclude dead cells from analysis. Cells were gated on FVD ^-^ population and analyzed on the BD FACS Aria III cytometer (BD Biosciences).

### Cytokine ELISA

DCs were pre-treated or not with EVs alone for 24 h and/or subsequently infected with *S. pneumoniae*, T4R for 2 h, followed by removal of extracellular bacteria and incubation for another 24 h. The culture supernatants were analyzed for cytokines using the Human TNF-α and Human IL-12 DuoSet ELISA kits (R&D Systems) following the manufacturer’s protocol.

### Zebrafish maintenance and EV microinjection

The zebrafish study was approved by the Institutional animal ethical committee, Indian Institute of Science, Bangalore, India (Ref. CAF/ETHICS/965/2023). Adult healthy wild-type zebrafish (*Danio rerio)* were procured from local vendors in Bangalore, India. The fish were maintained in DBGL Zebrafish Facility Centre, Indian Institute of Science, Bangalore, with continuous aeration with 14 h light/10 h darkness photoperiod and maintained at 28 ± 1 °C. The viable eggs were kept in 1X embryo medium 5.03 mM NaCl, 0.17 mM KCl, 0.33 mM CaCl_2_.2H_2_O, 0.33 mM MgSO4.7H_2_O) at 28.5°C. All microinjection procedures were carried out on 3 day post-fertilization embryos with the help of FemtoJet Microinjector and PatchMan NP2 Micromanipulator (Eppendorf, Germany). Embryos were divided into three groups-mock, Naïve EVs (EV), and PLY(0.5) EVs. 60 embryos were injected into the duct of Cuvier with 3 nl of the EV suspension at a consistent injection time of 0.02 sec and 90 hectopascal pressure. Following the microinjection process, embryos were sorted under a fluorescence microscope (Olympus BX51) and only fluorescence-positive embryos were transferred into the embryo medium and kept in a 28.5°C incubator with constant changing of media at 12-hour intervals.

### Zebrafish heartbeat, survival and real-time PCR

Zebrafish embryos were anesthetized using 0.04% Tricaine (Sigma-Aldrich, E10521) for experimental procedures. Sedated embryos were placed over an agarose mold and heartbeats were assessed manually using an Olympus SZ51 stereomicroscope. Zebrafish embryos were checked every day, dead larvae were removed and the fraction of surviving larvae was calculated. The experiment was performed in triplicates. The error bars represent the standard deviation of the survival percentage from the three independent experiments. Zebrafish embryos were imaged for four days post-injection using Zeiss 880-Multiphoton confocal microscope. Embryos were anesthetized as stated previously and agar mounted and expression for fluorescence-tagged EVs was measured in three microinjected groups of embryos. Images were processed using ImageJ software. qPCR experiments were performed using cDNAs prepared from the zebrafish embryos. Amplification of inflammatory cytokine TNF-α (Forward Primer: 5’-GGAGAGTTGCCTTTACCGCT-3’, Reverse Primer: 5’-GATTGCCCTGGGTCTTATGGAG-3’) was carried out with gene-specific exon-exon junction spanning primers using SOLIS BIODYNE HOT FIREPol EvaGreen kit following the manufacturer’s protocol. Fluorescence intensities were recorded and analyzed using Qiagen QIAquant 96 5-plex detection machine. Relative mRNA expression was determined by delta delta Ct method, taking into consideration the quantity of target sequences relative to the endogenous control β-actin (Forward Primer: 5’-CACTGAGGCTCCCCTGAATCCC-3’, Reverse Primer: 5’-CGTACAGAGAGAGCACAGCCTGG-3’).

### Mouse model of PLY-induced lung injury and adoptive transfer of EVs

All animal experiments were approved by the institutional animal ethics committee (Ref. IAEC/896/KARTHIKS/2022). 6-8 weeks-old wild-type C57BL/6 mice (50% male/female) were anaesthetized by inhalation of 4% isofluorane and 50□µl of PBS containing 0.25 μg of recombinant PLY (5 μg/ml) was instilled intranasally into the nostrils. For IVIS imaging experiments, GFP-tagged PLY was administered. To assess lung permeability, 50 μl of 0.5% Evans blue (Sigma) in saline was injected intravenously through the tail vein at 30 min prior to PLY administration as described previously (*51*). At 18 h post treatment, mice were sacrificed followed by heart perfusion to drain out blood and the lungs were imaged *ex vivo* for Evans blue dye lung extravasation (Excitation- 620 nm, Emission- 680 nm) and PLY-GFP (Excitation- 500 nm, Emission- 520 nm) signals in the lungs respectively using the IVIS Spectrum-CT Imaging system (Caliper-Perkin Elmer). Post sacrifice, lungs were perfused with PBS containing 2% FBS and 1 mM EDTA to collect the BALF. The BALF was processed to isolate the cells upon RBC lysis and stained with PE anti-mouse Ly6G clone 1A8 (Biolegend) and FITC anti-mouse F4/80 clone BM8 (Biolegend) for quantification of neutrophils and macrophages by flow cytometry (BD FACS Aria III). The EVs isolated from BALF of PLY and PBS treated mice were adoptively transferred to recipient mouse intranasally to recipient mice upon normalizing the total EV protein content to 300 μg/mouse. At 24 h, the mice were scored for clinical symptoms and the distribution of Nile-red EVs in mice was imaged using the IVIS system (Excitation- 535 nm, Emission- 620 nm). The infiltration of neutrophils and macrophages in BALF was quantified by flow cytometry. The supernatant of BALF was store at -80°C for cytokine analysis by mouse TNF-α and IL-1β DuoSet ELISA kits (R&D Systems). Lungs were perfused with a 1:1 mixture of OCT and 4% PFA and preserved in OCT cryomolds (Tissue-Tek) at -80°C. 10 µm sections were cryosectioned using the cryotome (CM1850UV Leica) and embedded in poly□l□lysine coated slides for H&E staining. Images were acquired through Olympus CKX53 Microscope under the 10x objective.

### Statistical analysis

Data were statistically analysed using GraphPad Prism v.9.1.2. Data represent mean±SEM unless otherwise specified. Comparison between multiple groups was performed using one-way ANOVA followed by Tukey’s multiple comparison tests as indicated. Normalized data were analysed using unpaired t-tests. Differences were considered significant at *P□<□0.05, **P□< 0.01; *** P<0.0005, NS, denotes not significant.

## Acknowledgements

The study was supported by extramural funding through the INSPIRE Faculty fellowship Department of Science and Technology (DST/INSPIRE/04/2019/002238), Start-up Grant (SRG/2021/000401) from the Science and Engineering Research Board (SERB), Ramalingaswami Re-entry fellowship (BT/RLF/Re-entry/46/2020) from the Department of Biotechnology, India and intramural funding from Rajiv Gandhi Centre for Biotechnology, Thiruvananthapuram. The authors profusely thank Prof. Birgitta Henriques Normark, Karolinska Institutet, Stockholm for sharing the pneumococcal strains used in the study. We extensively thank the veterinary medical officers, Dr. Arya Aravind and Dr. Archana at the animal house, RGCB for technical assistance with mouse experiments. We thank Dr. K.C Sivakumar, Bioinformatics facility, RGCB for the proteomics data analysis. The technical assistance provided by Mrs. Bindu Asokan, Mrs. Rintu T Varghese at the TEM facility, Mrs. Viji S at histology and Mrs. Surabhi S V at the Bioimaging facility, RGCB are also acknowledged.

## Author Contributions

K.S. conceptualized the project. K.S., S.P., C.V.B., S.D., S.S., Kuldeep. S., J.B.J., U.N., A.B., developed the methodology. K.S., J.B.J., U.N., A.B., acquired resources. K.S., C.V.B, S.A., A.J., A.D., S.D., S.S., Kuldeep. S., J.B.J., conducted the investigations. K.S., S.P., C.V.B., A.J., S.A., S.D., S.S., J.B.J., U.N., analysed the data. K.S. wrote the original draft. K.S., J.B.J., U.N., A.B., reviewed and edited the manuscript. K.S., J.B.J., U.N., A.B., acquired funding. K.S., J.B.J., U.N., A.B., supervised the project. All authors have reviewed and approved the manuscript.

## Data and materials availability

All data needed to evaluate the conclusions in the paper are present in the paper and/or the Supplementary Materials.

