## Supplemental figures for "Bacterial pore-forming toxin pneumolysin drives pathogenicity through shed toxin-loaded host extracellular vesicles"

**Supplementary Figures**

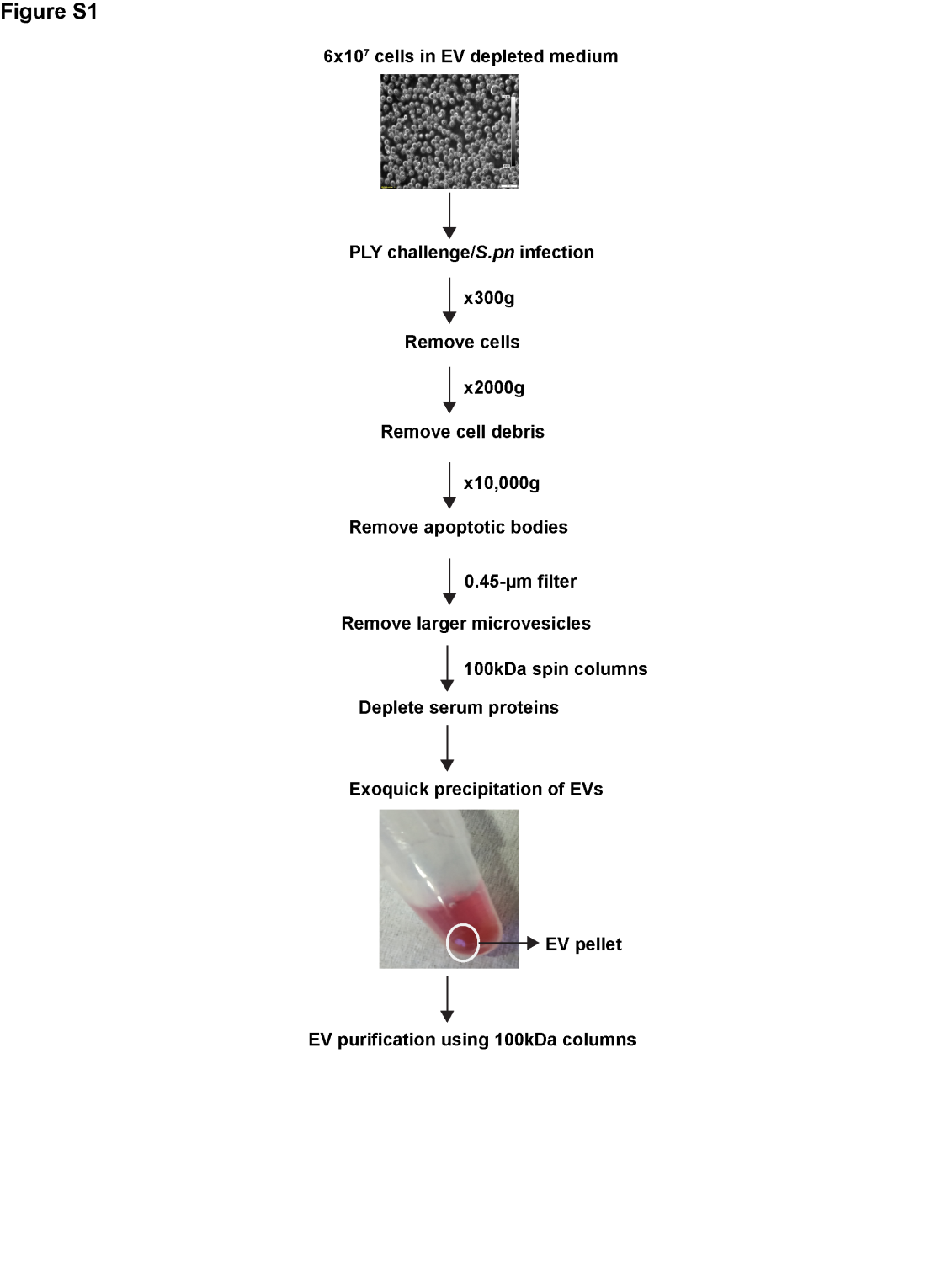

**Figure S1. Schematic representation of EV isolation procedure.** Briefly, 6x10^7^ cells were stimulated in culture medium depleted of serum EVs, following which differential centrifugation was used to remove cells, debris and apoptotic bodies. The conditioned supernatant was filtered through 0.45 µm syringe filters to remove larger microvesicles. The supernatant was concentrated using 100 kDa centrifugal spin columns to deplete non-EV proteins and the vesicles were subsequently precipitated using Exoquick reagent. The precipitated EVs were again washed and purified using 100 kDa centrifugal spin columns.

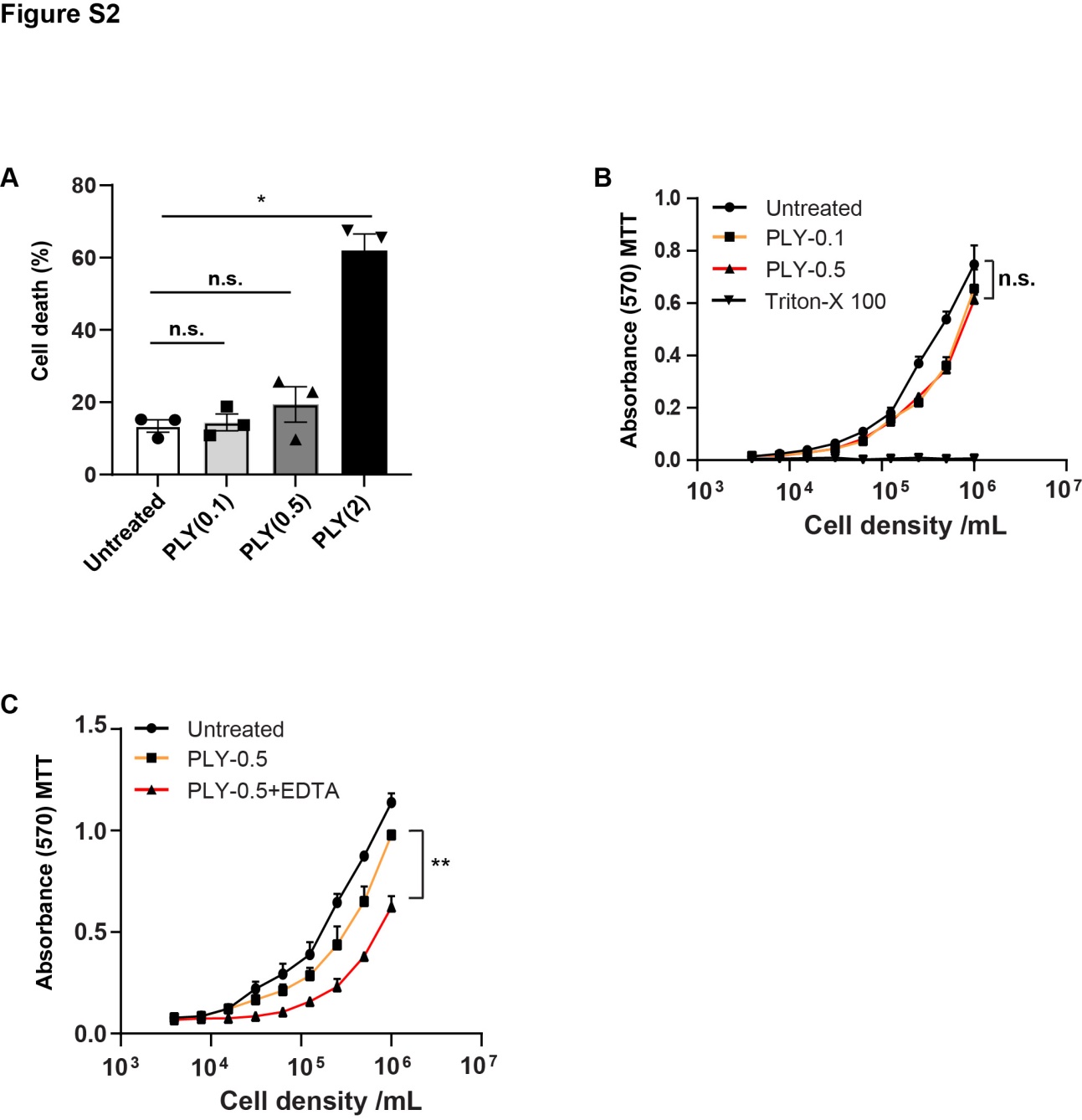

**Figure S2. Viability of THP-1 monocytes challenged with sublytic doses of pneumolysin** **(A)**. Cell death of THP-1 monocytes challenged with purified recombinant pneumolysin (0.1- 2 µg/ml) was measured by flow cytometric quantification of cells positively stained with the viability dye, FVD-efluor 780 (N=3). (**B**). MTT assay showing the viability of THP-1 monocytes treated with purified recombinant pneumolysin (0.1, 0.5 µg/ml). Treatment with detergent, Triton X-100 was used as positive control for cell death. (**C**). MTT assay showing the viability of THP-1 monocytes treated with purified recombinant pneumolysin (0.5 µg/ml) for 24h in the presence or absence of 25 mM EDTA. ** denotes p<0.005 by Wilcoxon matched pairs signed rank test. n.s. denotes not significant.

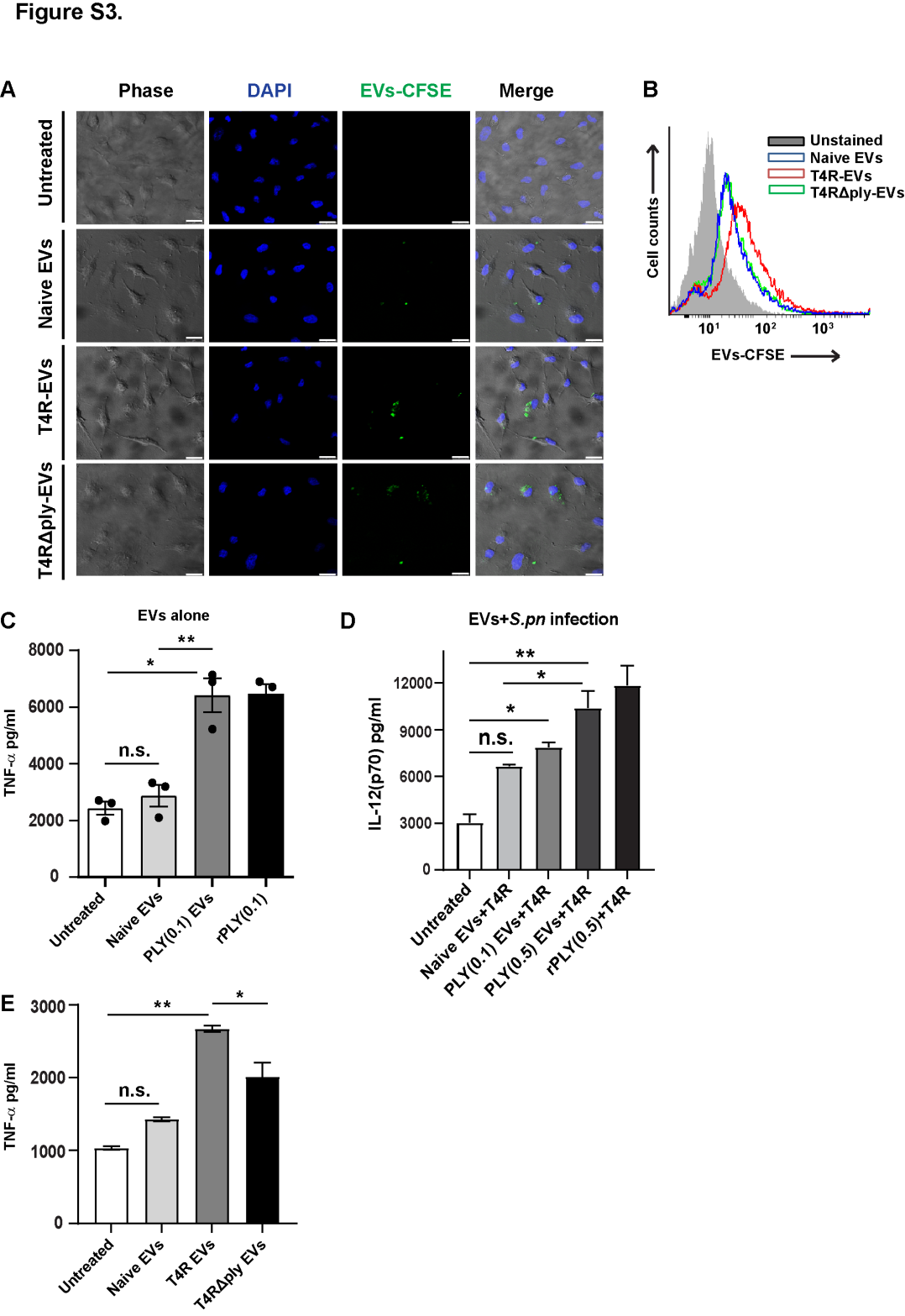

**Figure S3. EVs shedded from pneumococcal infected cells are internalized by alveolar epithelial cells and induce pro-inflammatory cytokine release. (A)**. Confocal microscopy showing the internalization of CFSE-labelled EVs (green) by human alveolar epithelial cells, A549 at 24h. The EVs were isolated from THP-1 monocytes infected with PLY-expressing unencapsulated serotype 4 strain, T4R (T4R-EVs), isogenic PLY mutant T4RΔPLY (T4RΔPLY-EVs) or uninfected cells (Naïve-EVs). Scale bars, 25 µm. (**B**). Flow cytometry histograms (N=2) to quantify the uptake of CFSE labelled T4R-EVs, T4RΔPLY-EVs and Naïve EVs by A549 cells at 24h post incubation. (**C-E**). Cytokine ELISA showing the levels of (**C**) secreted TNF-α from DCs treated with PLY (0.1)- or Naïve- EVs alone, (**D**) IL-12 from DCs pre-treated with PLY (0.1,0.5)- or Naïve- EVs for 24h followed by subsequent infection with *S. pneumoniae*, T4R strain and (**E**) TNF-α from DCs treated with T4R-EVs, T4RΔPLY-EVs or Naïve EVs. rPLY (0.5 µg/ml) was used as positive control. Data are representative of 2 independent experiments. * indicates p<0.05 and ** p<0.005 by one-way ANOVA with Tukey’s multiple comparisons test. n.s. denotes not significant.

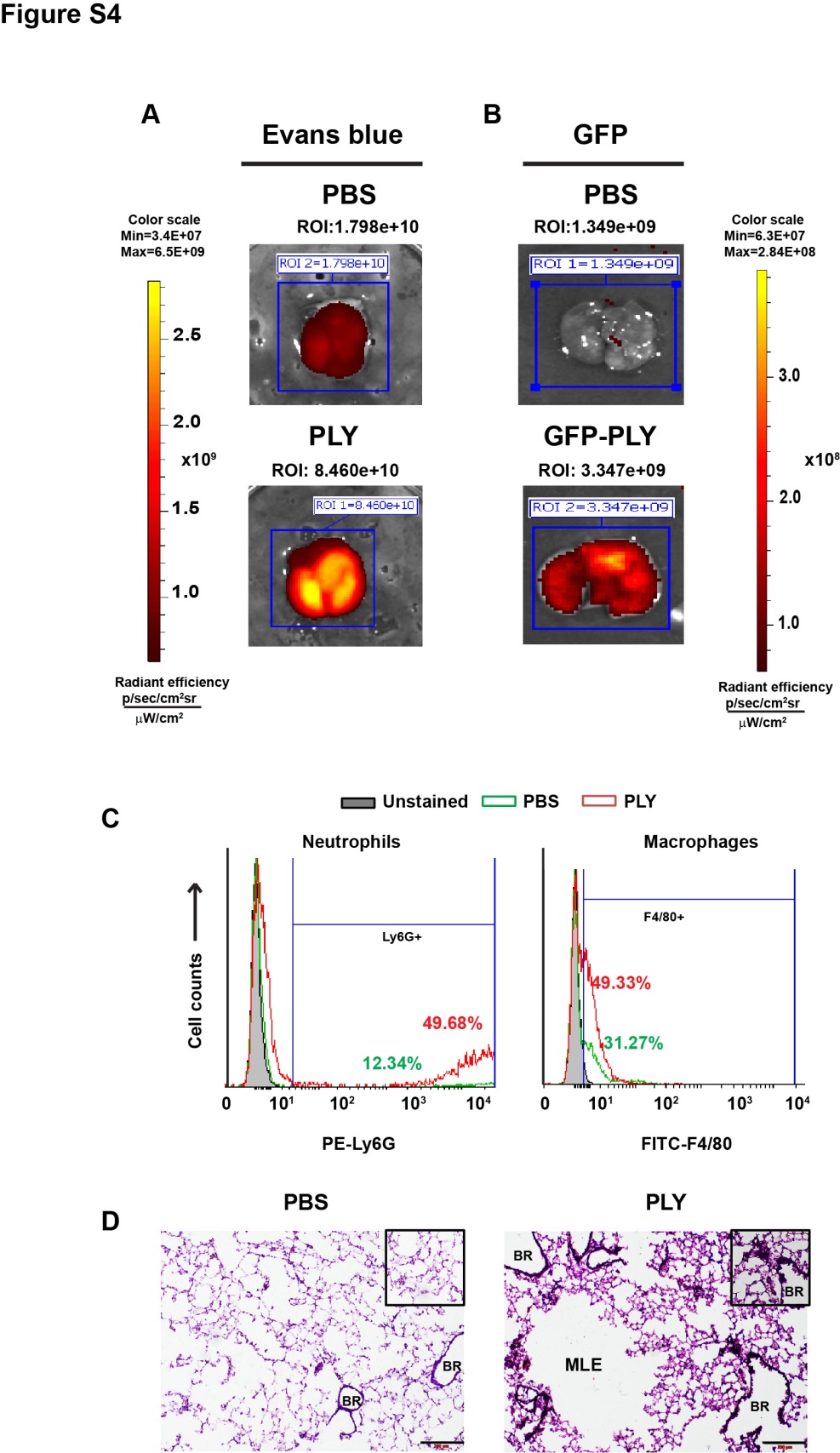

**Figure S4. Mouse model of pneumolysin induced acute lung injury. (A-B)**. IVIS imaging showing the lung distribution of (**A**) intravenously administered plasma tracer dye, Evans blue and (**B**) GFP-tagged PLY at 18h post intranasal administration of PLY (0.25 µg) or PBS into mice. The ROI intensity values are indicated showing higher Evans blue dye extravasation into lungs coinciding with higher PLY-GFP intensity. (**C**). Flow cytometry analysis of neutrophils (Ly6G^+^) and inflammatory macrophages (F4/80^+^) in BALF of mice treated with PBS or PLY (0.25 µg) at 18 h. (**D**). Hematoxylin and eosin (H&E) staining of mouse lungs at 18 h post intranasal instillation of PBS or PLY (0.25 µg). Mice administered with PLY showed tissue microlesions (MLE) and immune infiltration in the alveolar spaces confirming PLY-induced lung injury (magnified in the inset). BR-bronchiole; MLE-Microlesions. Scale bars, 200 µm.

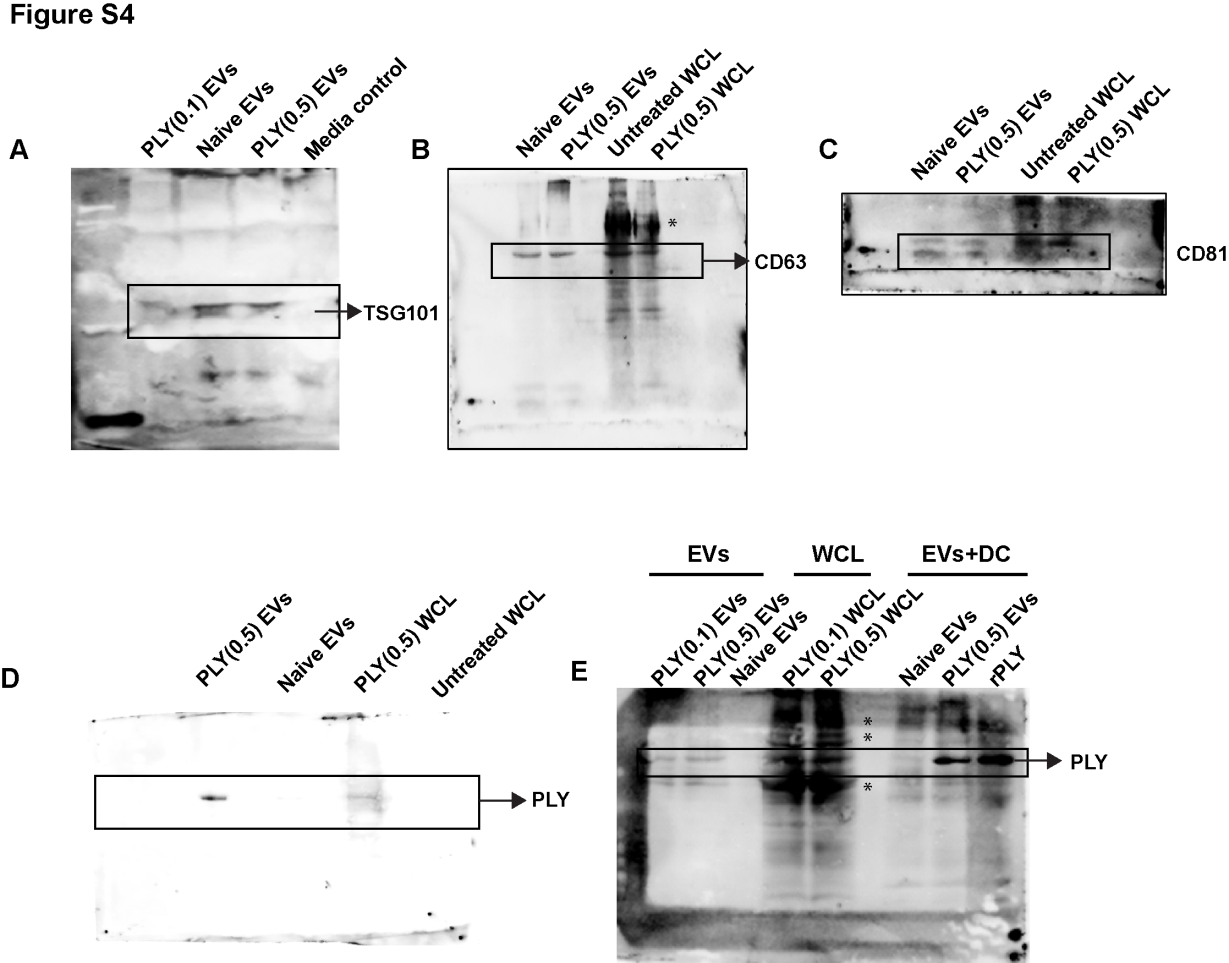

**Figure S5.** Original uncropped blots for Fig. 1E (panels A-C), Fig. 2A (panel D), Fig. 2C (panel E).* indicates non-specific band.

**Supplementary Videos:**

**Videos S1, S2. Pneumolysin upregulates EV shedding from THP-1 monocytes.** Human THP-1 monocytes were labelled with lipophilic membrane dye, Nile-red and stimulated with 0.5 µg/ml of purified recombinant PLY (**Video S1**) and PBS (**Video S2**) for 20 min and EV shedding was visualized by high-speed imaging at 5s intervals for a total of 20 min under the 63x oil objective of Leica SP8 laser confocal microscope. Videos are representative of 3 independent experiments. Scale bars, 10µm.

**Videos S3, S4**. **Pneumolysin is expelled from challenged THP-1 monocytes through shedded EVs**. Human THP-1 monocytes were labelled with lipophilic membrane dye, Vybrant Dil and stimulated with 0.4 µg/ml of recombinant GFP-tagged PLY (green) (**Video S3**) and PBS (**Video S4**) for 20 min and EV shedding was visualized by high-speed imaging at 5s intervals for a total of 20 min under the 63x oil objective of Leica SP8 laser confocal microscope. Videos are representative of 3 independent experiments. Scale bars, 25µm.
